# Effective Representation Learning via The Integrated Self-Supervised Pre-training models of StyleGAN2-ADA and DINO for Colonoscopy Images

**DOI:** 10.1101/2022.06.15.496360

**Authors:** Jong-Yeup Kim, Gayrat Tangriberganov, Woochul Jung, Dae Sung Kim, Hoon Sup Koo, Suehyun Lee, Sun Moon Kim

**Author notes:** Corresponding author: Sun Moon Kim.

## Abstract

In order to reach better performance in visual representation learning from image or video dataset, huge amount of annotated data are on demand. However, collecting and annotating large-scale datasets are costly and time-consuming tasks. Especially, in a domain like medical, it is hard to access patient images because of the privacy concerns and also not knowing what exactly to annotate without expert effort. One of the solutions to obviate the hassle is to use Self-Supervised learning methods (SSL) and Generative Adversarial Networks (GANs) together. SSL and GANs are quickly advancing fields. GANs have unique capability to create unlabeled data sources containing photo-realistic images while SSL methods are able to learn general image and video features from large-scale data without using any human-annotated labels. In this work, we explore leveraging the power of recently introduced StyleGAN2-ada and Self-Supervised Pre-Training of Dino together for the pretext task. Our underlying insight is also that leveraging the current approaches with Transfer Learning (TF) together brings benefit on doing pretext task in medical domain. By the strategy of unifying these two approaches, we propose the integrated version and use it derive representation learning on polyp dataset.

## Introduction

Artificial intelligence (AI) already powerfully assists human beings in the effective performance of countless tasks. Applications with built-in deep learning have become ubiquitous. Familiar examples include recommendation systems that help customers make quick and wise decisions, and self-driving cars that use powerful AI algorithms to reduce the likelihood of accidents [1, 2]. Physicians will undoubtedly rely on AI in the near future. Image-based medical diagnostic specialties, such as pathology, radiology, and endoscopy, are expected to be the first medical fields to be affected by the introduction of convolutional neural networks [3]. Deep learning for lesion classification and detection is expected to help endoscopists provide more accurate diagnoses.

Nevertheless, almost all deep learning related projects can achieve good performance due to the datasets which have annotation data. On top of that, some domains have a data insufficiency problem. For instance, medical domain not only faces less possibility of finding patient images but also a difficulty of image labelling issue. In fact, first problem is a lack of enough dataset due to the privacy while second one is annotating medical that requires time and many doctors efforts to label object on medical images. Thus, we refer to Self-supervised learning (SSL) approaches not requiring any annotation for image representation learning. To tackle the insufficiency of medical data, we leverage Generative Adversarial Network (GAN) supplying enough dataset for our target.

GAN itself is a model for image generation to reconstruct a similar image to input image. GANs working structure is that two networks compete with each other. The discriminator network competes with the generator and tries to distinguish between the realistic fake images coming from the generator, which samples z vector from a latent space to generate, and the real samples. GANs learn to create realistic images that are similar to the input images but not present in the input dataset. Following the tic-toe game approach, the discriminator and generator are enhancing their abilities simultaneously. This is because generator enforces to learn how to generate realistic images according to discriminator’s level of distinguishing power that is arising empirically during the train [4].

The idea behind this is that if GAN can supply a large realistic polyp dataset for instance, then we must have solved an insufficient data problem noticeably. From this point, it can be the key to privacy problems that yields an alternative to medical image collection. StyleGAN2-ada [5] is one of the outstanding generative models that can provide various realistic images with the limited input data.

Generally, SSL pipeline derives pretext and downstream tasks. In pretext task, feature extraction can be done while downstream task includes object detection or segmentation. Mostly, the pretext task part of SSL refers to Vision Transformers (ViT) related models.

Vision Transformers (ViT)s [6] have brought big changes and turned into new trend in computer vision [7] and new investigation for medical data [8] after inspiration by natural language processing (NLP), that is, pre-training on large quantities of data and fine-tuning on the target dataset [9]. Transformers have unusual capacity for demonstration in learning pretext tasks. Especially, they show their efficiency when extracting global and local representations through layers, and provide scalability for large-scale training. As an alternative to convolutional neural networks (CNNs) having restricted receptive fields, ViTs encode visual features from a set of patches and use self-attention blocks in order to model long-range global information [10]. However, ViTs have some drawbacks, which cause them to lag behind from CNN architectures, such as a demand on huge amount of training dataset, and their extracted features are sensitive in exhibiting properties. Recently there have been some works to improve ViT performance. One of them is DINO [11] model composing ViT with knowledge distillation.

Taken the viewpoint of the vital role of pretext task in SSL into consideration, in this paper, we focus on how to get an effective representation learning in pretext task. Because 95% of success on implementation depends on pretext task. In addition, Transfer-Learning assists to boost the performance during the process.

Since the efficacy of SSL can rely on the dataset generated by GAN, we bridge Self-supervised visual representation learning and GAN. The integrated models with transfer-learning setup are complementary, *i.e.* leading to strong performance on the pretext tasks.

Our two-step performances can be summarized as follows:

- We generated fake polyp images (300k) using one of the contemporary GAN version such as StyleGAN2-ada with pre-trained model and vice verse (no pre-trained model).
- We applied two generated polyp datasets to Self-Supervised Pre-Training of Dino models trained on ImageNet and No Pre-Training of Dino models for pretext task respectively.

Since GAN itself is referred to as a Generative Self-Supervised Learning that does not require labeled data for training, we can see hybrid combination of two types of SSL architecture in the current demonstration.

## Related works

### Generative Adversarial Network

(GAN) has started revolutionary trend in computer vision due to the GAN-generated images that have been extensively utilized as a source for training models [12, 13, 14]. GANs are now capable of producing highly realistic images that are tough to distinguish when comparing with the data on which they are trained. For instance, StyleGAN [15] has an inversion of finding the latent code of real images in the latent domain. It has been used by many prior works [16, 17] as prevalent image synthesis model due to its latent space semantic richness.

However, StyleGAN just shows good performance when big data to assure the sufficient training of the discriminator is available. Later on, Karras et al. [5] proposed StyleGAN2-ada in which Dynamic data-augmentation strategy tailored for StyleGAN2 for the best training on few-sample datasets. StyleGAN2-ADA makes StyleGAN applicable to tasks with limited data by using adaptive discriminator augmentations. StyleGAN2-ada also focuses on distinct facets such as synthesizing qualitative images with high resolution and supporting various levels of style control. To this end, we found StyleGAN2-ada high quality and diverse image generator for providing to our training model with sufficient training data.

### Self-Supervised Learning

(SSL) [18] refers large-scale representation learning through self-supervision without the demand of data annotation by human intervention in comparison to Supervised counterparts. SSL has pretext as well as downstream tasks. First task is about feature representation learning from unlabelled dataset while second ones includes dense prediction tasks, e.g., image classification, object detection and image segmentation. Pretext task mainly relies on ViT models for feature extractions in SSL.

### ViTs

is a subsequent improvement of Transformers. After Transformers [19] gained initial achievements in NLP [20, 21], There have been continuous efforts on generalizing Transformers to computer vision [22, 23]. This remarkably reduces the architectural gap between NLP and vision.

### DINO

[11] approach itself dates back to Transformers by ViTs. To be exact, Vision Transformer (ViT) [6] was in turn proposed to adapt the Transformer for image recognition by tokenizing and flattening images into a sequence of tokens.

ViTs are representative image encoders being composed of multi-head attention blocks. DINO is an extended version of ViTs that adopt knowledge distillation and designing an unlabeled self-supervised method with knowledge distillation by analyzing the characteristics of ViT features.

In medical context, For image and video analysis problems, a number of pretext tasks have been explored, including prediction of image rotation [24], position prediction [25], colorization [26] and image context restoration [27, 28] etc.

To precise, Tajbakhsh et al. [24] chose image rotation for the pretext task and providing lung lobe segmentation and nodule detection tasks with the useful pretrained features in the downstream task. Bai W et al. [25] formulated anatomical position prediction as the pretext task for cardiac MR image segmentation Ross et al. [26] used colorization as pretext task and further applying the extracted features to initialize a surgical instrument segmentation model. Le Thi et al. [27] accomplished image reconstruction as pretext task on polyps segmentation task while Dippel [28] et al. extends contrastive loss to a self-reconstruction task with attention mechanism on fundus images.

DINO also is a type of pretext task toolkit. Yet, to emerge distinct properties of medical images like polyps, the network requires a large unlabeled corpus of images. In fact, our observation on polyp dataset reveals that models pretrained on DINO suffer performance drop when we used transfer learning with KvasirSEG in low data regime. Moreover, polyp dataset is one of the datasets, which tends to be noisy for foreground and background separation. It might be partly attributed to learn feature representations and increasing the complexity for further image analysis. Transfer Learning can mitigate this issue.

### Transfer Learning

ImageNet pre-training is the de-facto standard practice for many visual recognition tasks [29]. The weights pre-trained on ImageNet is used to initialize the Transformer backbone for training on downstream tasks.

However, we demonstrate the transferability of DINO pre-trained models on synthetic polyp dataset for pretext task not downstream task. It implies that Self-supervised DINO ViT backbones trained on ImageNet with no labels are not helpful with the small datasets in medical domain. To deal with the present issue we explore that if it is fine-tuned through DINO weights for our current experiment using huge polyp dataset, then it performs favourably. To prepare a vast number of polyp data points, StyleGAN2-ADA pre-trained model on the FFHQ 1024 × 1024 dataset is trained using an off-the-shelf dataset (KvasirSEG).

## Materials and Methods

### The Kvasir-SEG

[30] is an open source polyps dataset specialized for Deep Learning-relevant demonstrations e.g., polyps detection, localization, and segmentation tasks. The high-resolution electromagnetic imaging system such as ScopeGuide, Olympus Europe acquired 1000 polyp images. The resolution of the images in this dataset ranges from 332 × 487 to 1920 × 1072 pixels. The dataset is available online at https://datasets.simula.no/kvasir-seg/. The dataset includes and 48 small polyps (≤ 64 × 64 pixels).

### GANs

were tough to train with small dataset, often facing mode-collapse, where the discriminator starts to memorize all original images, resulting in the overfitting and no longer guiding the generator well. There has been recent advance such as **StyleGAN2-ADA** [5] to avoid this behaviour by augmenting the dataset to prevent its memorization. StyleGAN2-ADA proposes an adaptive discriminator augmentation mechanism to maintain stability of training with limited data. The augmentation mechanism consisting of groups of image transformations (e.g., cropping, resizing) with the probability values is applied to both real and fake images. In particular, the augmentation pipeline includes 18 transformations, as well as an adaptive hyper-parameter to control the strength of these augmentations.

### DINO

[11] relies on teacher-student strategy.

- **In theoretical perspective**, DINO performs self-distillation on two different augmented views of the same image using a similar teacher-student setup where discards the dissimilar representations by training a student network to predict the representations generated by the student network that is an exponential moving average teacher network.

*x* – input image,
g_*θs*_, g_*θt*_ – student and teacher networks,
*θ_s_*, *θ_t_* – networks parameters,
*P_s_*, *P_t_* – probability distributions (outputs of the networks) respectively. *K* – output probability distributions of the networks over dimensions denoted by *P_s_* and *P_t_*.
*T_s_*, *T_t_* – temperature parameters (*T_s_*, *T_t_* > 0).

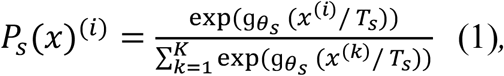

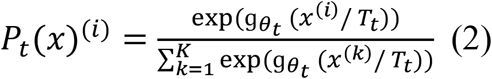 The equation (1) holds true with the equation (2) for probability distributions (*P_s_* and *P_t_*).
- **In practical perspective**, DINO demonstrate that self-supervised ViTs can automatically highlight salient foreground objects with the power of the produced attention maps, even though they were trained with no supervision as well as their features explicitly contain the scene layout, object boundaries including the information regarding semantic location of an image.

We adopted a two-stage approach for our pretext task: A former stage is to have a sufficient synthetic dataset by training StyleGAN2-ADA pre-trained model on small polyp dataset (Kvasir-SEG). A latter stage is to fuel DINO pre-trained models with the synthetic dataset.

## Experiments and Results

In this section we describe details of our implementation. Our experiment has two stages including generating fake images for training setup and training DINO with the generated fake images. We trained StyleGAN2-ADA using KvasirSEG dataset and generated 300K fake images. This huge fake dataset acts as training-set to train DINO for pretext task. We also did extra small implementation for downstream task to have quantitative output in the end.

We trained StyleGAN2-ada with pre-trained and no pre-trained sets. For the evaluation on StyleGAN2-ada, Fréchet Inception Distance (FID) is calculated. FID is a metric to evaluate GANs. FID captures the similarity of generated images to real ones better than Inception Score. Table 1 shows that StyleGAN2-ada with pre-trained mode is better that no pre-trained one. We generated 300K fake images twice (pre-training/no pre-training) for DINO training setup. We have used default sets of hyperparameters for the optimization of DINO models unless otherwise noted. The same scenario has repeated for DINO. To be more exact, we trained DINO with pre-training/no pre-training. Figure 2 show qualitative image comparison on Kvasir-SEG dataset. Figure 3 illustrates qualitative image comparison on Konyang dataset for both pre-training/no pre-training sets. We have found that the transfer learning performance of DINO manages to surpass the performance of DINO that was trained without transfer learning.

**Fig 1:**
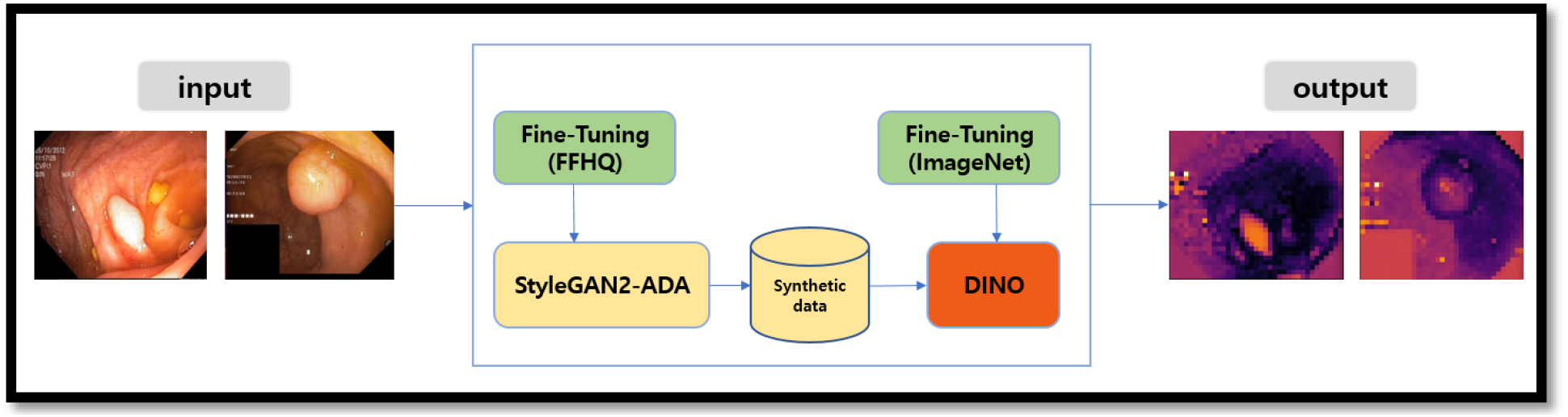
two-stage SSL approach for pretext task

**Fig 2:**
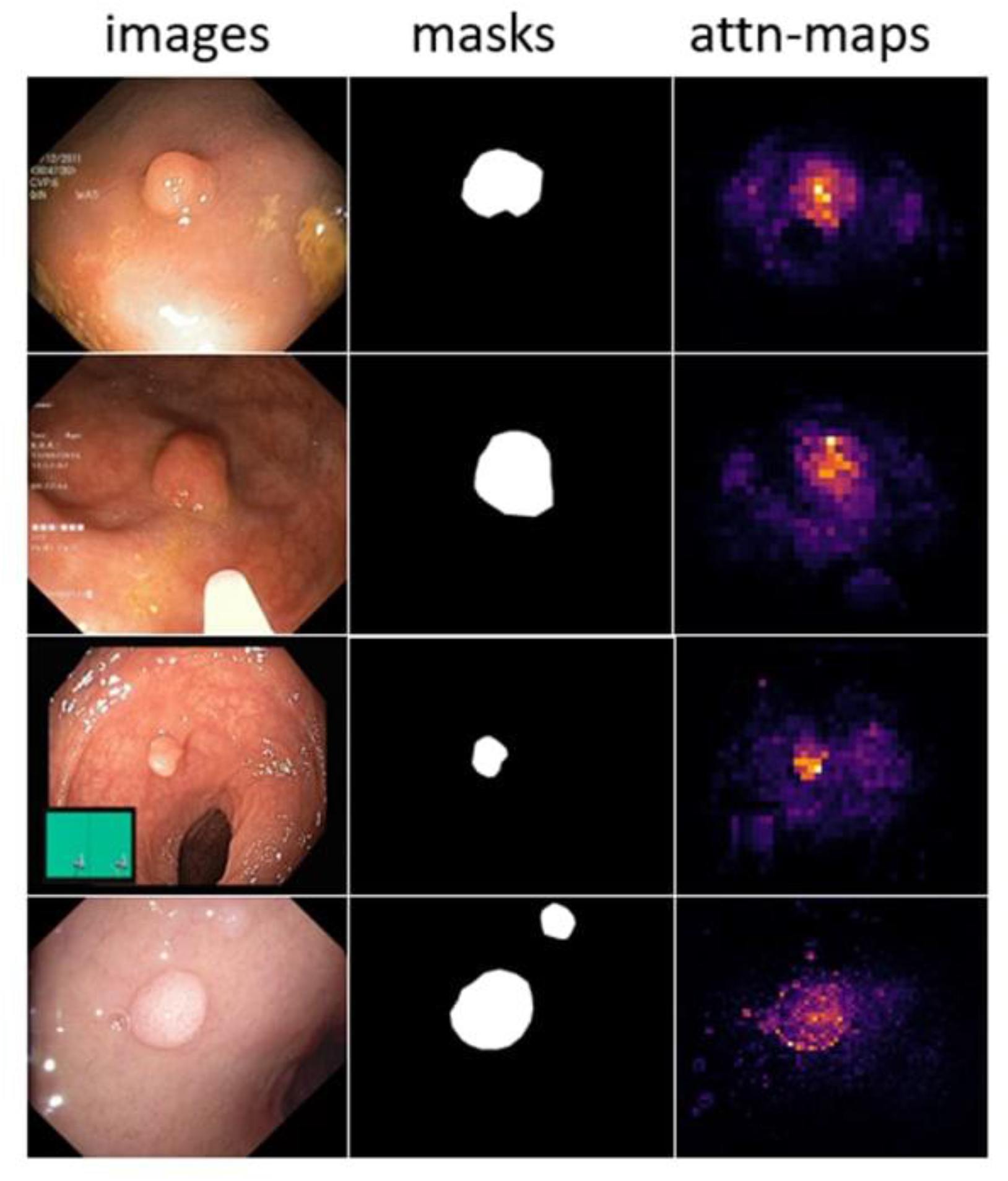
Qualitative results comparison on the Kvasir-SEG dataset. From the left: image (1), (2) Ground truth, (3) Dino attn-maps outputs.

**Fig 3:**
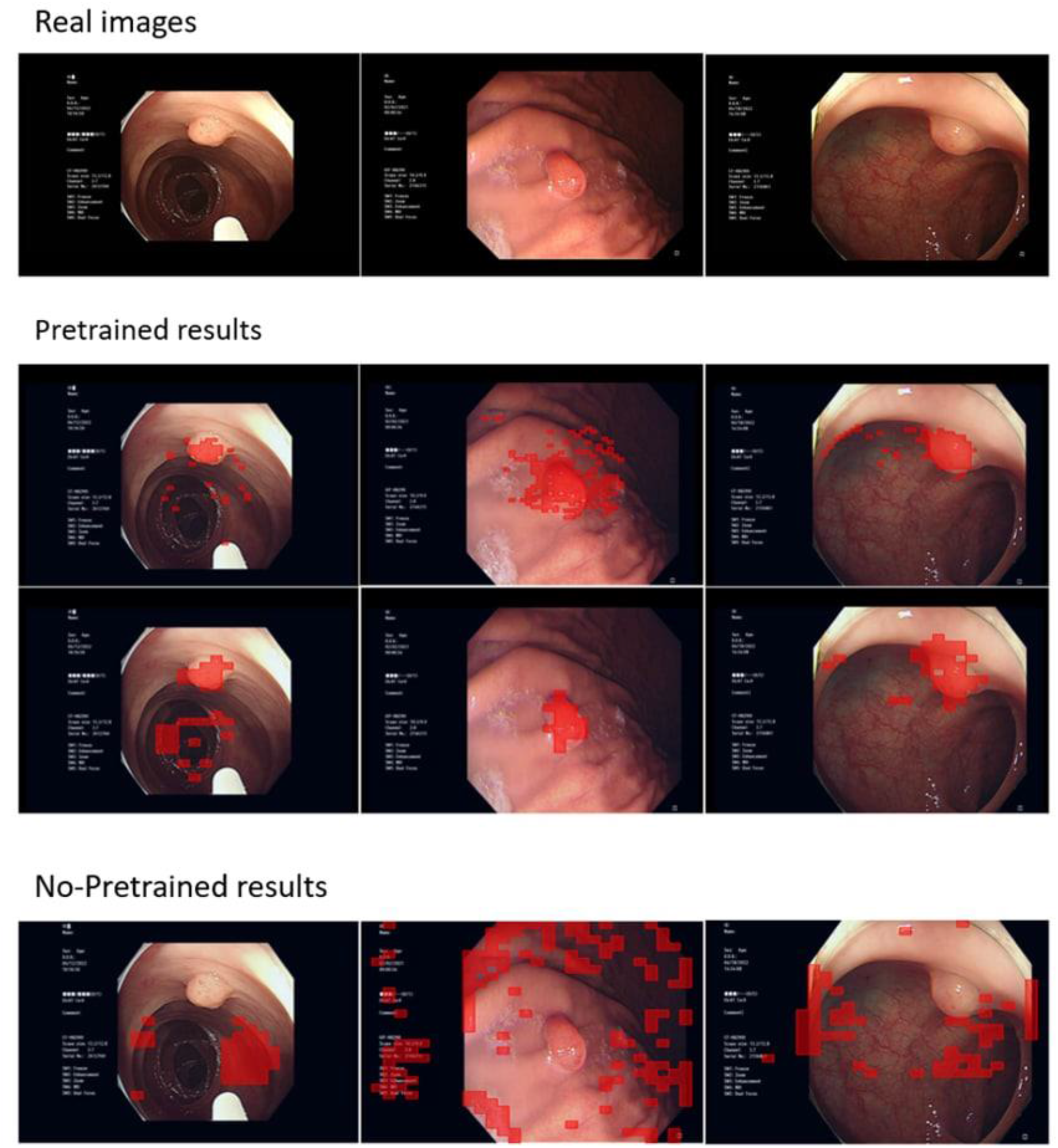
We visualize masks obtained by thresholding the self-attention maps. We show the resulting masks for a ViTs trained with DINO. We show the heads for both pre-training and no pre-training sets.

**Table 1:** KvasirSEG polyp dataset trained with StyleGAN2-ADA, from scratch and using transfer learning. We used FFHQ-140K with matching resolution as a starting point for all transfers. We compute FID by following the way of Heusel et al. [31].

| Model | FID |  |
| --- | --- | --- |
|  | No Pre-training | Pre-training |
| StyleGAN2-ADA | 31.57 | 26.02 |

The best heads can help to improve the performances in downstream tasks, e.g., polyp segmentation [32]. For instance, we made small dataset using Konyang video frames with no labels. We named it as selected data. The selected data contain 100 polyp images with appropriate masks. The masks have been prepared based on the best head samples from DINO through post-processing. Table 2 and Figure 5 shows recall, precision, dice coefficient, mIoU scores and ROC curves respectively for two datasets including Kvasir-SEG and Kvasir-SEG with selected data. The merge of Kvasir-SEG and selected data can boost performance on polyp segmentation. However, we conjecture that the selected data caused a decrease of mIoU score. The selected data masks are slightly inaccurate. Because they have been prepared based on DINO attention head maps not based on expert knowledge.

**Fig.4:**
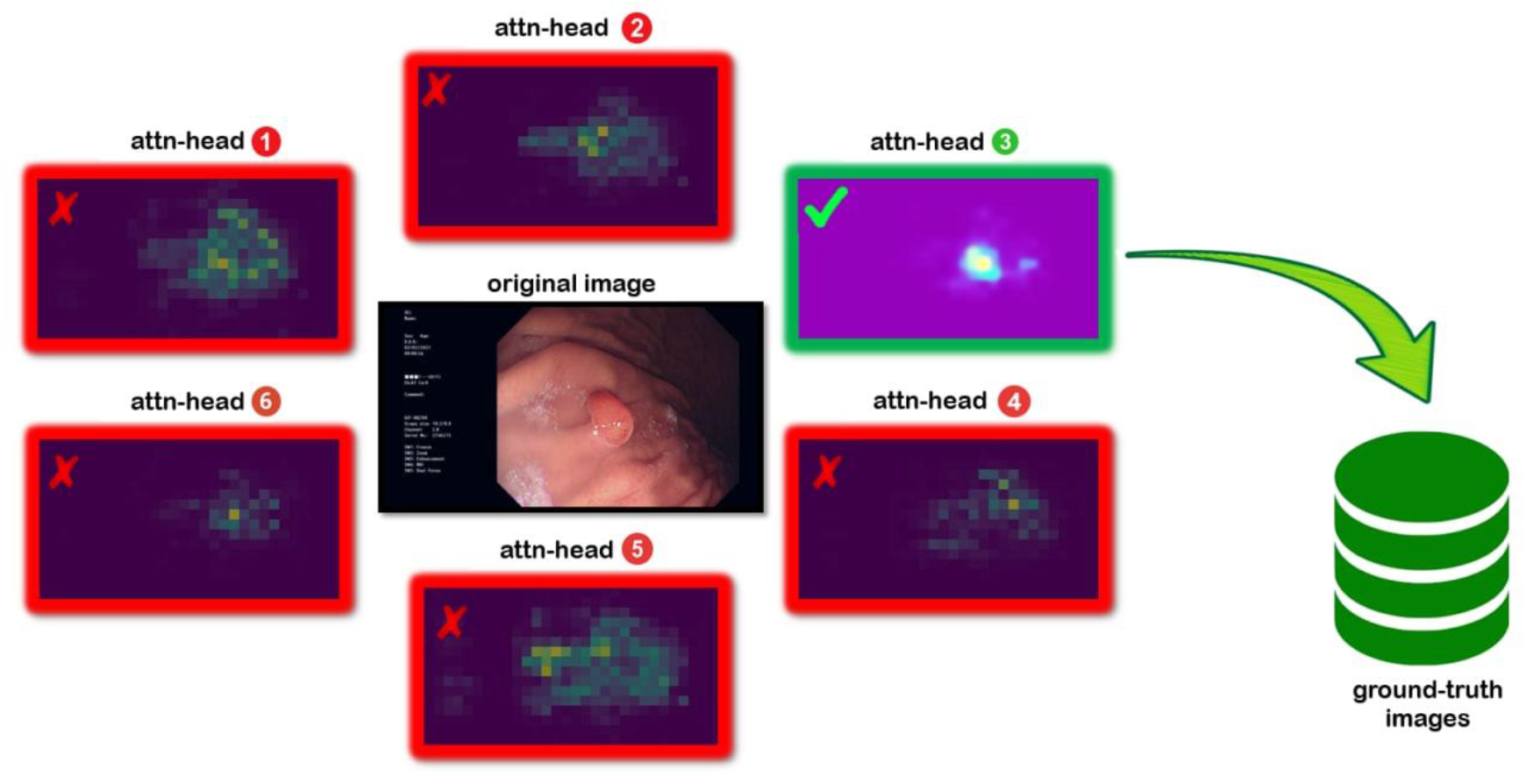
We show the best head selection in the post-processing.

**Fig.5:**
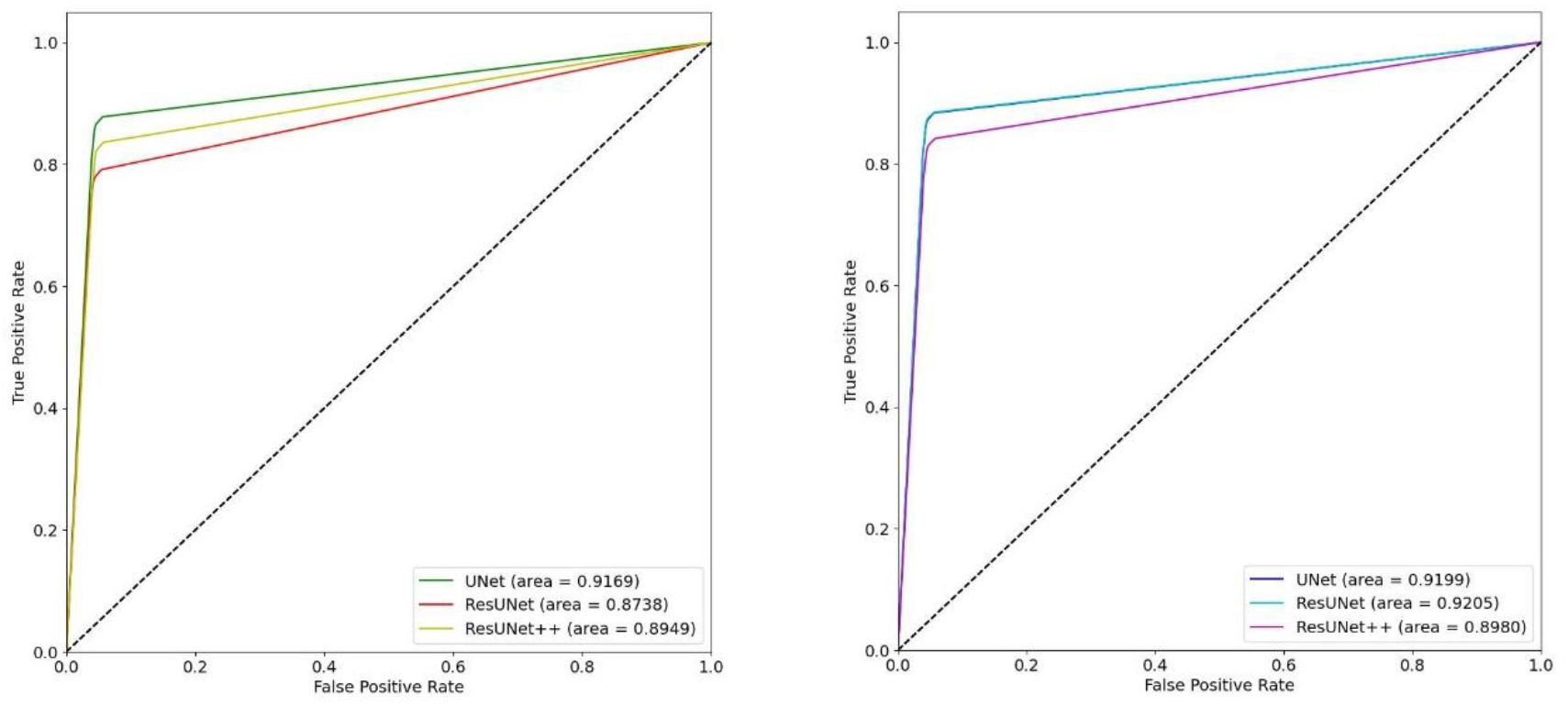
ROC Curves results. From the left: (1) Kvasir-SEG, (2) selected_data +Kvasir-SEG

**Table.2:**
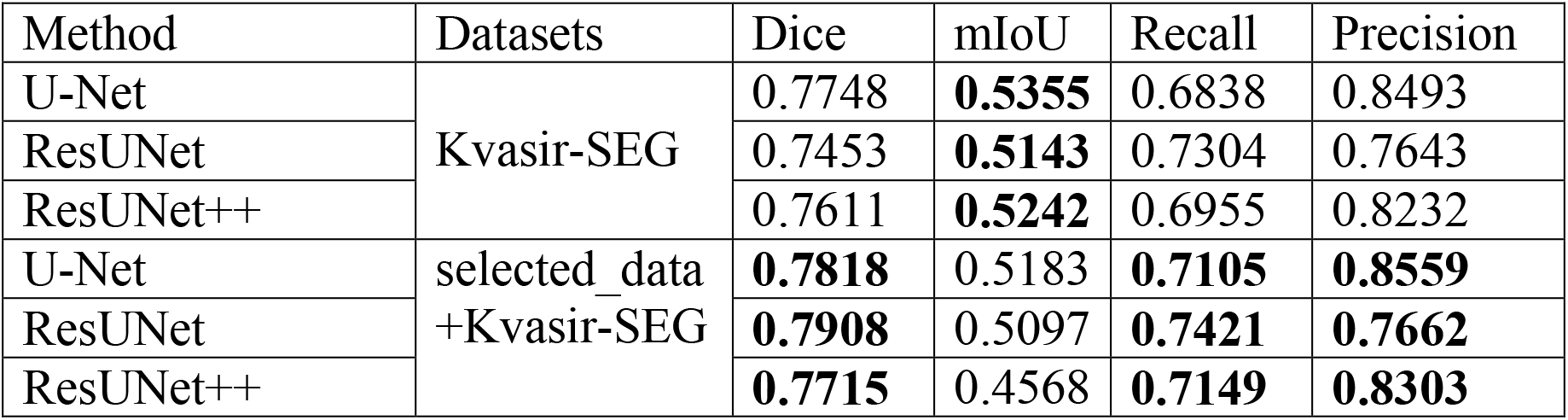
The quantitative results for two datasets.

| Method | Datasets | Dice | mIoU | Recall | Precision |
| --- | --- | --- | --- | --- | --- |
| U-Net | Kvasir-SEG | 0.7748 | **0.5355** | 0.6838 | 0.8493 |
| ResUNet |  | 0.7453 | **0.5143** | 0.7304 | 0.7643 |
| ResUNet++ |  | 0.7611 | **0.5242** | 0.6955 | 0.8232 |
| U-Net | selected_data<br>+Kvasir-SEG | **0.7818** | 0.5183 | **0.7105** | **0.8559** |
| ResUNet |  | **0.7908** | 0.5097 | **0.7421** | **0.7662** |
| ResUNet++ |  | **0.7715** | 0.4568 | **0.7149** | **0.8303** |

## Discussion

### GAN

In medical domain, researchers face two main issues including data collection and labelling. Firstly, it is known that the exploit medical data on any purpose with no patient’s agreement leads to break patient’s privacy (intellectual property rights). Secondly, data labelling is a laborious and costly venture in that Doctors collaborations are in need. However, synthetic dataset [33] also circumvents some of the concerns around privacy and usage rights that limit the spread of real datasets. To this end, GANs has emerged as a promising solution since they provide a data source so that further enables us train our models. We opted to some fake and real image samples for getting a cosine similarity value for which we provide examples in Figure 6. Cosine similarity is adopted for similarity measurement. Given a query image from synthetic polyp image set, it measures the probability of any correct matching in real polyp images (with same category label). It yields 87.6% similarity in between real and synthetic polyp images.

**Fig.6:**
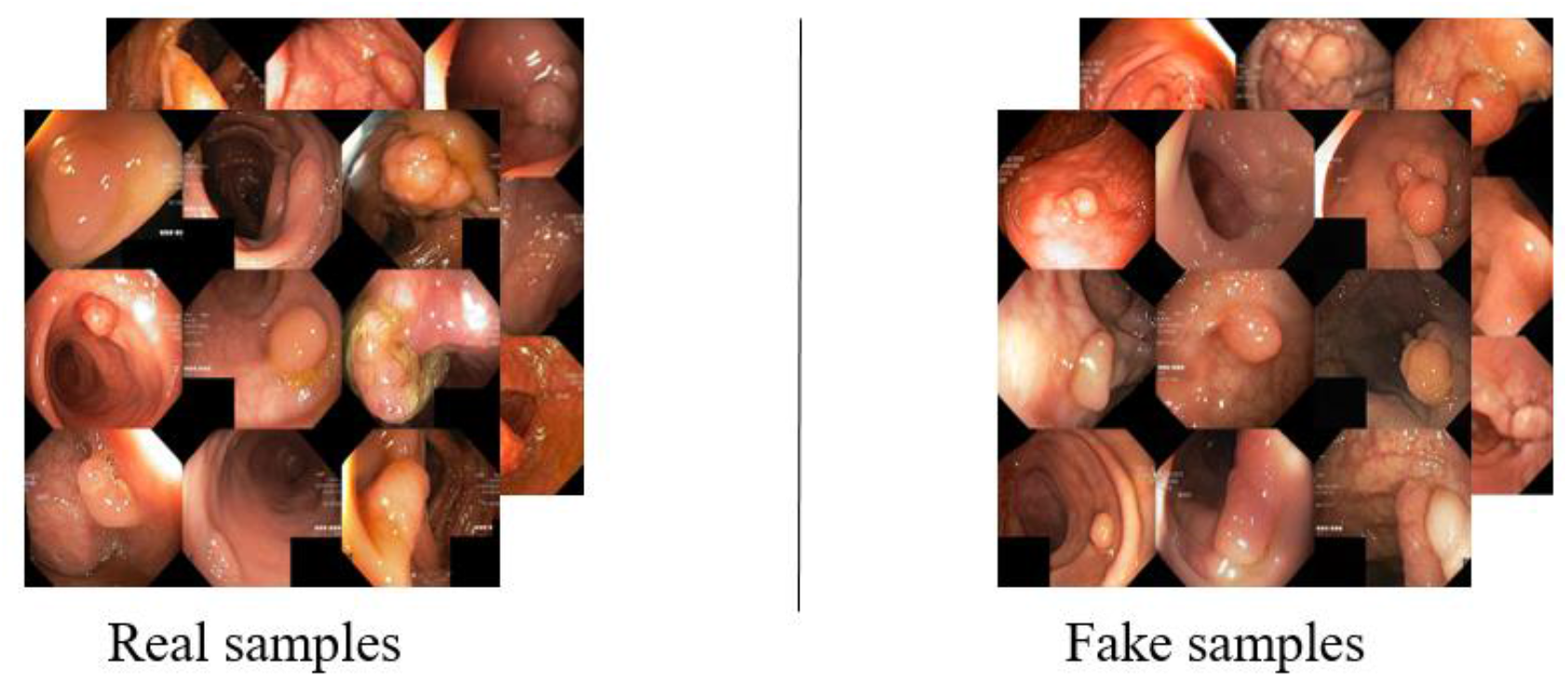
shows real image samples and fake image samples

### DINO

Typically, deep learning pre-trained models are fine-tuned with a small dataset. However, we decided to prepare a huge dataset (300K) for our feature extraction task for two reasons below.

Firstly, as illustrated in Figure 6, there is an issue about polyp image composition. To be exact, foreground images and background images are mostly in the same colours, which makes images cluttered. As a result, model tends to fail in producing high-quality attention maps in such ambiguous situation. This also impedes image-level and pixel-level representation learning in pretext task.

Secondly, unlike Convolutional Neural Networks (CNNs), DINO architectures do not inherently encode inductive biases (prior knowledge) to deal with visual data. They typically require large amount of training to figure out the main modality-specific rules. For example, a CNN has inbuilt translation invariance, weight sharing, and partial scale invariance due to pooling operations or multi-scale processing blocks. However, a Transformer network needs to figure out these image-specific concepts on its own from the training examples. Similarly, relationships between video frames need to be discovered automatically by the self-attention mechanism by looking at a large database of video sequences. This results in longer training times, a significant increase in computational requirements, and large datasets for processing. For example, the ViT [6] model achieved state-of-the-art on ImageNet classification by directly applying Transformers with global self-attention to full-sized images in that requires hundreds of millions of image examples to obtain reasonable performance on the ImageNet benchmark dataset. The question of learning a Transformer in a data-efficient manner is an open research problem and recent works report encouraging steps towards its resolution. For example, DeiT [34] uses a distillation approach to achieve data efficiency while T2T (Tokens-to-Token) ViT [35] models local structure by combining spatially close tokens together, thus leading to competitive performance when trained only on ImageNet from scratch (without pre-training). Another approach to data efficient training of ViTs is proposed in et al. [36]. The authors show that by smoothing the local loss surface using sharpness-aware minimizer (SAM) [36], ViTs can be trained with simple data augmentation (random crop, and horizontal lip) [38], instead of employing compute intensive augmentation strategies. All the mentioned recent Vision Transformers demonstrate that the model can be learned end-to-end on ImageNet-1K without any dedicated pre-training phase. Yet, in medical domain (e.g., polyp dataset), it performs favourably when experiment utility derives a huge polyp dataset from which pre-trained Dino’s multi-head attention modules learn to segment salient foreground objects as a byproduct of solving the pretext task.

### Transfer learning

As stated in [5], transfer learning provides significantly better results than from-scratch training. Hence, instead of random initialization, the initializing with a StyleGAN2-ADA model pre-trained on the FFHQ 1024 × 1024 dataset can speed up convergence and reduce data requirements. The same way adopted to pretext task in that DINO weights pre-trained on ImageNet are utilized to initialize ViTs on StyleGAN2-ADA-generated polyps dataset. In addition, the experiment is done without transfer learning mode to comparison.

## Conclusion

In this work, we observe that DINO tends to automatically learns class-specific attention maps leading to foreground object segmentations, even though its query located in foreground or background are in similar colours. But, a large dataset is highly required for training set as DINO architectures are hungry for data. Hence, StyleGAN2-ADA is able to serve as an adequate data supplier. Transfer Learning also shows more benefits than without pre-training mode.

## Acknowledgement

The Institutional Review Board of Konyang University Hospital (KYUH) approved this study and waived the requirement for informed consent, considering the retrospective study design and the use of anonymized patient data.

